# Convection and extracellular matrix binding control interstitial transport of extracellular vesicles

**DOI:** 10.1101/2022.08.03.502657

**Authors:** Peter A. Sariano, Rachel R. Mizenko, Venktesh S. Shirure, Abigail Brandt, Cem Nesiri, Bhupinder Shergill, David M. Rocke, Randy P. Carney, Steven C. George

**Author notes:** contributed equally as co-senior authors (Randy P. Carney, Steven C. George). Corresponding Author: Steven C. George, M.D., Ph.D., Professor and Chair, Department of Biomedical Engineering 451 E. Health Sciences Drive, room 2315, University of California, Davis, Davis, CA 95616.

## Abstract

Extracellular vesicles (EVs) influence a host of normal and pathophysiological processes *in vivo*. Compared to soluble mediators, EVs are relatively large (~30-150 nm) and can traffic a wide range of proteins on their surface including extracellular matrix (ECM) binding proteins. We isolated EVs from the MCF10 series – a model human cell line of breast cancer progression – and demonstrated increasing presence of laminin-binding integrins α3β1 and α6β1 on the EVs as the malignant potential of the MCF10 cells increased. Transport of the EVs within a microfluidic device under controlled physiological interstitial flow (0.15-0.75 μm/s) demonstrated that convection was the dominant mechanism of transport. Binding of the EVs to the ECM enhanced the spatial concentration and gradient, which was partially mitigated by blocking integrins α3β1 and α6β1. Our studies demonstrate that convection and ECM binding are the dominant mechanisms controlling EV interstitial transport and should be leveraged in the design of nanotherapeutics.

## Main

Over the past decade it has become clear that small (~30-150 nm diameter) extracellular vesicles (EVs) represent a distinct and fundamental component of how neighboring and distant cells and tissues communicate. The breadth of observations is impressive, including normal biological processes, such as cell migration ^1,2^, differentiation ^3,4^, and proliferation ^5,6^, as well as pathological processes such as priming the metastatic niche in cancer ^7^. EVs are composite nanoparticles, secreted by cells, that are comprised of a lipid-based membrane surrounding an aqueous core. The membrane and core can each incorporate a wide range of biologically-active molecules (e.g., proteins, nucleic acids). These characteristics endow EVs with numerous desirable qualities (e.g. stability in circulating biofluids ^8^; relatively rapid cellular uptake ^9^) which might be leveraged in the design of nanotherapeutics. Despite the enormous potential of EVs as a nanotechnology, there are currently no FDA-approved EV formulations, and our fundamental understanding of their endogenous biology remains in its infancy.

While much work has been dedicated to characterizing the function of EVs in the tissue microenvironment, much less is known about their physical location, distribution, and concentration within the interstitium. The interstitial space broadly describes the tissue microenvironment between the blood and lymphatic microvasculature and is characterized by a porous extracellular matrix (ECM) and interstitial flow (fluid exits the capillary bed and is reabsorbed by blood capillaries and lymphatics). Pore size is heterogeneous and depends on the tissue, but there is significant evidence, largely from electron microscopy in adipose, skin, and breast tissue, to suggest a wide range (50-5000 nm) capable of allowing relatively uninhibited transport of EVs ^10–13^. EVs are resident constituents of the ECM ^14,15^, and can influence ECM architecture and structure through aggrecanases, matrix metalloproteinases, elastase, and likely other matrix digesting proteases ^16–19^. EVs also have the potential to be transported through the ECM by diffusion ^20^ and convection, but also actively bind to collagen, laminin, and fibronectin via adhesion molecules (e.g., integrins) present on the surface of EV populations ^14,19,21^.

Despite the fact that interstitial flow is a fundamental feature of essentially all tissue microenvironments, and can profoundly impact the spatial distribution of soluble mediators ^22,23^, its role in manipulating and transporting EVs within the interstitium has been largely unexplored. This is particularly curious given the relatively large size of EVs (relative to small molecules) and thus relatively low rates of diffusion (diffusivity on the order 10^-13^ m^2^/s in tissue-like hydrogels ^20^ compared to 10^-10^-10^-11^ m^2^/s for small molecules in tissue).

Here we characterized diffusion, convection, and binding of EVs derived from the MCF10 human breast cell line series in a microfluidic device under controlled flow conditions through a laminin-rich ECM. The MCF10 series (MCF10A, normal; MCF10DCIS, pre-malignant; MCF10CA1, malignant) is widely used in both *in vitro* and *in vivo* studies of breast cancer ^24–27^, including several studies assessing EVs isolated from these parent cells ^28–31^. Furthermore, there is relatively high expression of adhesion molecules on cancer derived EVs ^32^, and several proteomics studies have demonstrated increasing presence of relevant integrins on EVs from increasingly malignant parent epithelium, including integrins α3, α6, and β1 ^28,33^. We demonstrate from both experimental and computational methods that under physiologic flow conditions, convection (not diffusion) and binding control EV transport in a laminin-rich ECM; furthermore, concentration of bound EV and spatial gradients are exaggerated as malignant potential increases, and these observations are due, in part, to enhanced expression of integrins α3, α6, and β1.

### EV features during malignant progression

We first isolated and characterized EVs from each line of the MCF10 series using nanoparticle tracking analysis (NTA) and transmission electron microscopy. EV size distributions (**Fig. 1a**) were consistent with reported features, with most measured particles within the expected range <150 nm. EVs secreted per cell increased more than 2-fold as the malignant potential of the cell line increased (**Fig. 1b**). This result is consistent with observations demonstrating that EV secretion is often increased in cancerous cells due to signaling dysregulation ^34^ or microenvironmental factors such as acidic pH or hypoxia ^35^. EV size distribution was confirmed by TEM, with most demonstrating the characteristic cup-shaped morphology caused by dehydration and rupture of hydrated particles during TEM preparation (**Fig. 1c**).

**Figure 1.**
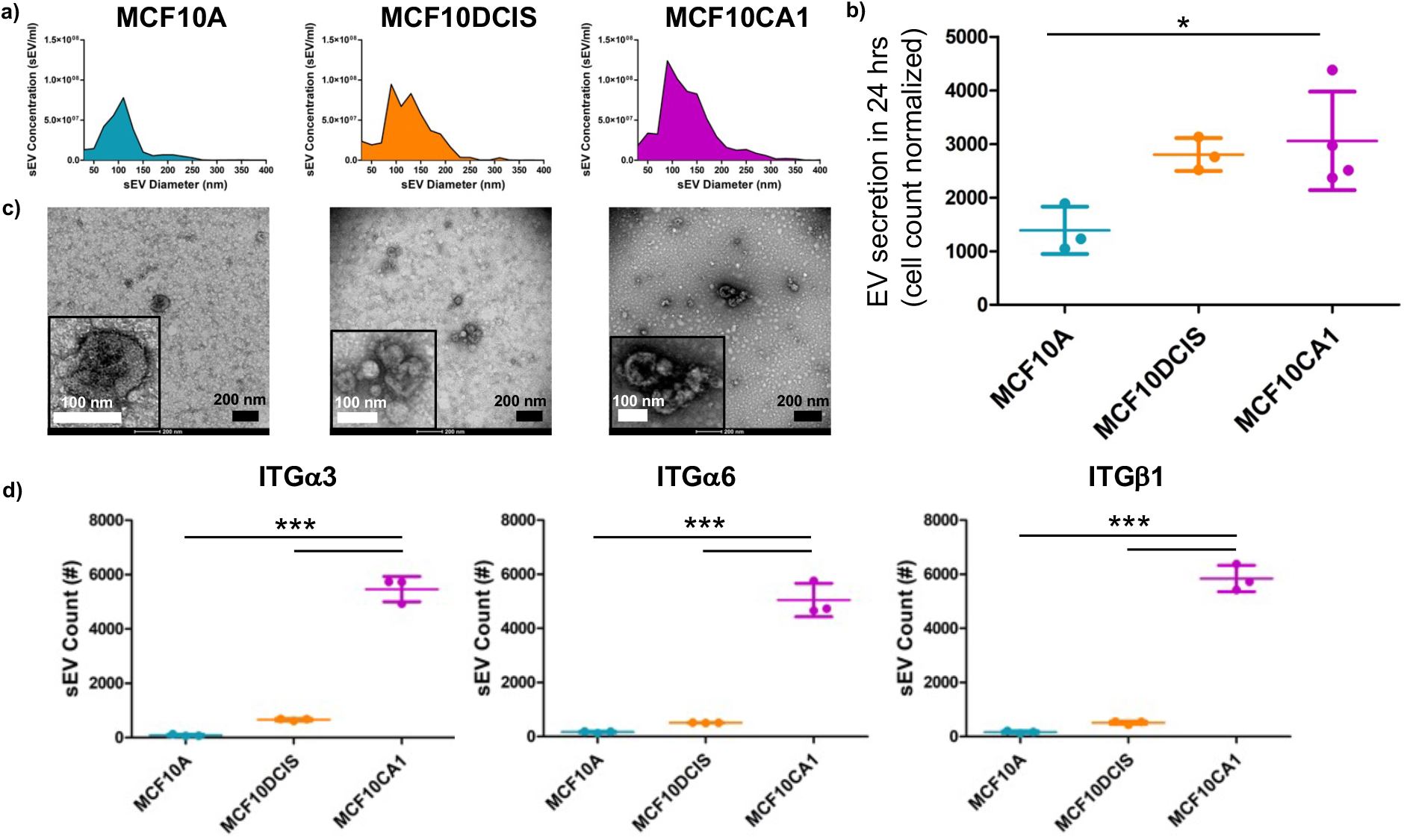
Isolation and characterization of EVs. a) Nanoparticle tracking analysis (NTA) revealed expected EV size distribution and adequate concentration for in vitro studies as well as b) increasing secretion with malignancy when normalized by cell number. c) Transmission electron microscopy (TEM) demonstrated sufficient EV concentration and a characteristic cup-shape morphology. d) Loading the same concentration of EVs revealed increasing expression of integrin α3, α6, and β1 with parent cell malignancy by ExoView analysis. *p=0.0319; ***p<0.0001; One-way Anova, Tukey post-hoc.

Using a dual antibody capture immunofluorescence technique (ExoView), we confirmed that EVs from all three cell lines express common tetraspanin markers (CD9, CD63, and CD81) enriched on cell-released EVs (**Fig. S1a**). We also observed significant EV heterogeneity in tetraspanin presence across the MCF10 series which supports the observation that EVs from distinct cell sources can carry markedly different protein profiles ^36,37^, and that these profiles can transform with malignancy.

We next assessed integrin expression on the EVs. EV samples isolated from each cell line were stained with a non-specific dye (Cell-trace Far Red, CTFR), immobilized (ExoView) and probed for laminin-binding integrins α3, α6, and β1. Integrin positive EVs were determined by thresholding against the MIgG negative control. Comparison across the lines revealed significantly increased expression of all three integrins on MCF10CA1 EVs compared to normal or pre-malignant MCF10A or MCF10DCIS EVs (**Fig. 1d**). There was no significant difference between MCF10A and MCF10DCIS EVs. Integrins require dimerization of α and β subunits to effectively bind ligand targets ^38^, which prompted analysis of the colocalization of integrins α3, α6, and β1 on individual EVs. The presence of colocalized integrins on individual EVs (**Fig. S1b,c**; cyan+gray fractions) increased 2-4-fold with increasing parent cell malignancy for both α3β1 and α6β1 integrin pairs (**Suppl. Table 1**).

### EV diffusive transport and binding

We next assessed the functional consequence of integrin expression on the EVs by characterizing the diffusivity and binding rates to a laminin-rich ECM using fluorescence recovery after photobleaching (FRAP). FRAP analysis affords the ability to not only measure EV diffusion in bulk (i.e., molecular diffusivity), but also to derive kinetic binding parameters including forward (k_on_), reverse (k_off_), and equilibrium K_d_ (k_off_/k_on_) rate constants. We modelled the process of ECM binding as a pseudo first order process, 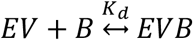, where B is the concentration of binding sites and is considered in excess. The kinetic parameters can be determined from the shape of the fluorescence recovery profile following photobleaching (**Fig. S2a**) using non-linear regression of a diffusion-reaction model ^39^. In addition to the diffusivity and kinetic binding parameters, the asymptotic level reached by FRAP curves is indicative of a bleached immobile fraction as well as the unbleached freely diffusing mobile fraction (**Fig. S2a**).

The FRAP methodology and the analysis pipeline were first validated by performing bleaching experiments with 40 kDa and 150 kDa dextran (**Fig. S2b,c**). We used a 3 mg/mL laminin-rich ECM (Matrigel-GF Reduced; Corning) that produces a matrix with comparable pore size range as that present *in vivo* ^40^. The fluorescent signal from the dextrans recovered to nearly baseline, thus reflecting the expected lack of binding to matrix (no immobile fraction). The diffusivity of the 40 kDa and 150 kDa dextran were extracted from the fitted recovery curves (1.6 x 10^-11^ m^2^/s and 0.94 x 10^-11^ m^2^/s, respectively), and were ~40% of the theoretically estimated (Stokes-Einstein) ^41^ diffusion coefficient in water (3.5 x 10^-11^ m^2^/s and 2.6 x 10^-11^ m^2^/s, respectively); thus consistent with diffusion in tissue-like hydrogels ^42^.

For integrins to form heterodimer pairs, divalent cations such as Ca^2+^ or Mg^2+^ are necessary to stabilize the interaction, and Mn^2+^ can be used to lock the integrins in a high affinity-binding state. As such, we modelled a low, intermediate, and high affinity integrin binding state of the EVs using either EDTA to chelate free cations or functionally-validated blocking antibodies (α3, α6, β1, and β4), CaCl_2_ + MgCl_2_, or MnCl_2_, respectively. The fluorescence recovery curves for each of the cell lines and treatment conditions were then fit for K*on (product of kon and the concentration of binding sites), K_d_, and D, then combined, averaged, and plotted with 95% confidence intervals. For each condition, EVs isolated from MCF10CA1 exhibited slower initial recoveries and a lower plateau, consistent with ECM binding (**Fig. 2a**). This observation was followed by MCF10DCIS EVs which shared 95% confidence intervals with MCF10A EVs at most time points.

**Figure 2.**
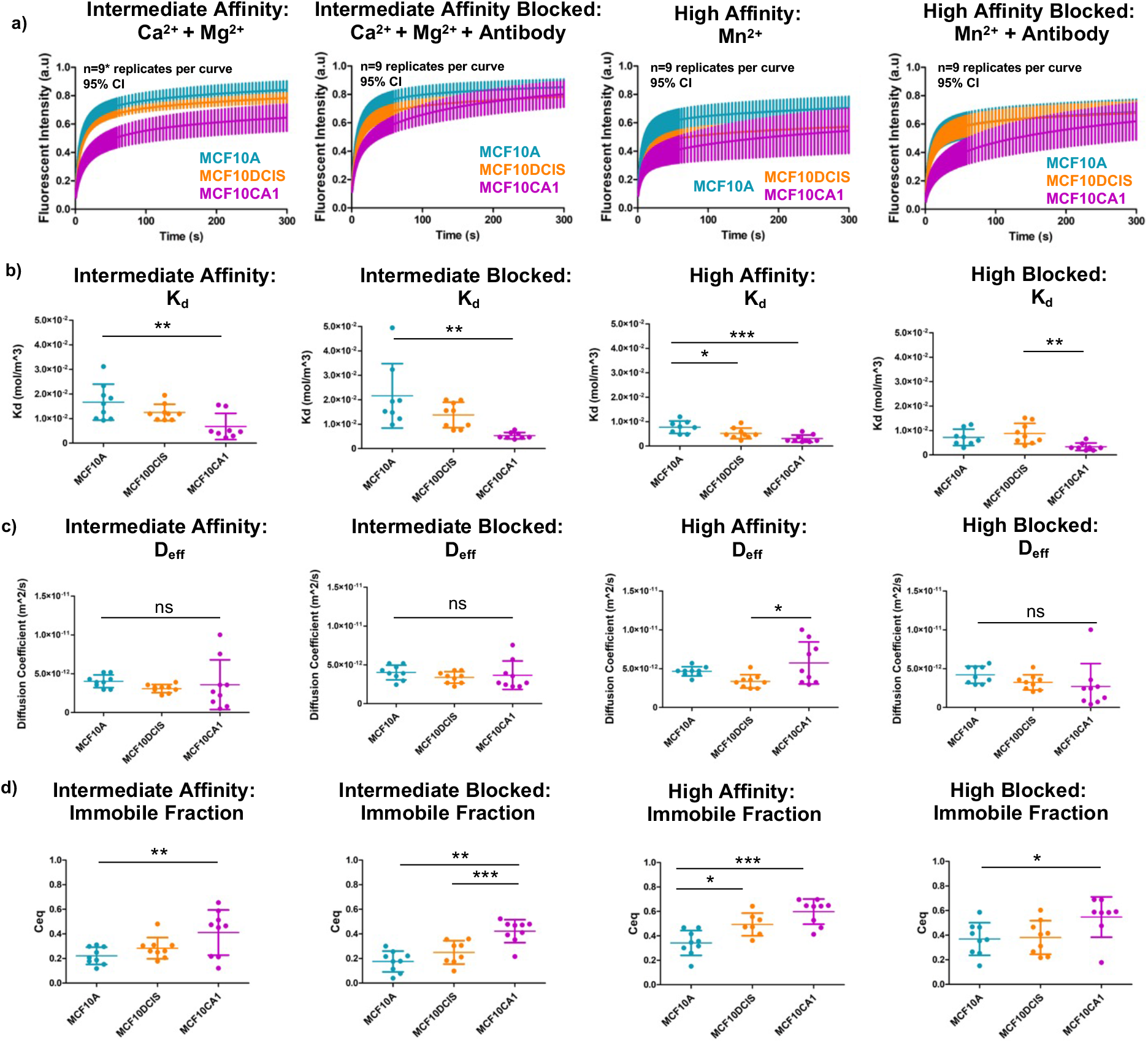
EV matrix binding is impacted by parent cell malignancy. a) FRAP recovery fitted curves were consistently slower and lower for malignant MCF10CA1 EVs across all treatment conditions, followed by MCF10DCIS and MCF10A EVs. Recoveries generally appeared to display asymptotic behavior by t= 300sec. Error bars= 95% CI; n=9 replicates per curve; n*=8 replicates for MCF10CA1 curve. b) Fitted K_d_ parameter was consistent with behavior in FRAP curves. MCF10CA1 EVs exhibited lowest K_d_ values, indicating the highest levels of EV binding to the matrix. c) Molecular diffusivity was broadly independent of cell line or treatment condition. d) The bound fraction (C_eq_) was assessed for each integrin binding state. Highest EV binding fractions were observed with MCF10CA1 EVs in all conditions compared to EVs from MCF10A and MCF10DCIS. One-way Anova, Tukey post-hoc. *p<0.05, **p<0.01, ***p<0.001; One-way Anova, Tukey post-hoc.

We next fitted for the values of k*_on_, K_d_, and D for each of the curves to better understand the mechanism leading to the altered fluorescence recovery curves. EVs from the MCF10CA1 line consistently exhibited the lowest K_d_ across all conditions (**Fig. 2b**). Integrin blocking had minimal impact on K_d_. In contrast, *D* was not impacted by cell line and integrin activation conditions with the exception of a small increase in MCF10CA1 in the high affinity condition (**Fig. 2c**). The mean value for *D* (3.80 x 10^-12^ m^2^/s) is consistent with previous estimates using nanoparticle tracking ^20^, and is within the range of theoretical values (3.4-17 x 10^-12^ m^2^/s) of a sphere in water with a mean diameter ranging from 30-150 nm. Comparing the immobile, bound EV fraction (C_eq_) for each condition (**Fig. 2d**) resulted in similar trends to K_d_. In other words, the increasing immobile fraction of EVs observed with increasing malignant potential is consistent with enhanced binding (lower K_d_) to the extracellular matrix. The enhanced binding could be due to either a reduced k_off_, an enhanced k_on_, or a combination. A near constant value of *D* for the EVs across cell lines is consistent with Stokes-Einstein theory and the characterization of the free diffusion of EVs from each cell line, which demonstrate a similar size distribution (**Fig. 1**).

### *In silico* model of EV convective transport and binding

With properties of EV diffusion and binding now characterized, we next investigated the role of physiological interstitial flow (convection) on the transport of EV in a laminin-rich ECM using a finite element computational model (COMSOL). The geometry of a previously described microfluidic device ^43,44^ was imported into COMSOL (**Fig. S4**). The microfluidic device consists of five parallel chambers that can communicate by diffusion and convection through a series of small evenly dispersed pores. The pressure distribution and porosity create a physiological interstitial flow from the central chamber outwards through hydrogels loaded in the adjacent compartments. All modeling parameters were gathered from the literature or determined experimentally through iterative *in vitro-in silico* experiments (**Suppl. Table 2**).

We first demonstrated that physiological flow (velocity, range 0.1-10 μm/s) could be achieved using three different hydrostatic head pressure heads (5.5, 8.8, and 12.5 mm). These pressure heads produced mean interstitial flow velocities of 0.5, 0.75. and 1 μm/s, respectively, along the centerline between opposing pores across the interstitial microfluidic channel (**Fig. 3a**). To assess the impact of convection on bound and free concentration of EVs, we introduced a constant concentration of EVs in the microfluidic line adjacent to the interstitial channel on the high-pressure side to allow EVs to enter the interstitial channel by convection. We considered no binding, intermediate (Kd of MCF10A in the presence of MgCl_2_ and CaCl_2_), and high (K_d_ of MCF10CA1 in presence of MnCl_2_) (**Suppl. Table 2**). Across all integrin binding states and initial binding site concentrations, occupancy of binding sites was negligible, consistent with the fact that the binding site concentration was calculated to be several orders of magnitude higher than EV influx concentration. EV accumulation in the interstitial compartment increased with increasing EV binding affinity (**Fig. 3b**). Concentration gradients between opposing pores in the interstitial channel were evident, but very small in magnitude, in both free and bound EV species (**Fig. 3c**). Faster interstitial flow velocities flattened free and bound gradients, particularly in the high affinity binding condition, consistent with our earlier experimental and theoretical observations of dextran and vascular endothelial growth factor ^43^ (**Fig. 3c**). Bound and free EV concentrations were similar in magnitude, although the bound fraction surpassed free EV concentrations in the high affinity binding condition. Further, high affinity binding resulted in nearly a two-fold increase in total EV concentration (free + bound concentration) compared to intermediate or no binding conditions. These observations highlight the potential biological significance of both free and bound fractions in the interstitium.

**Figure 3.**
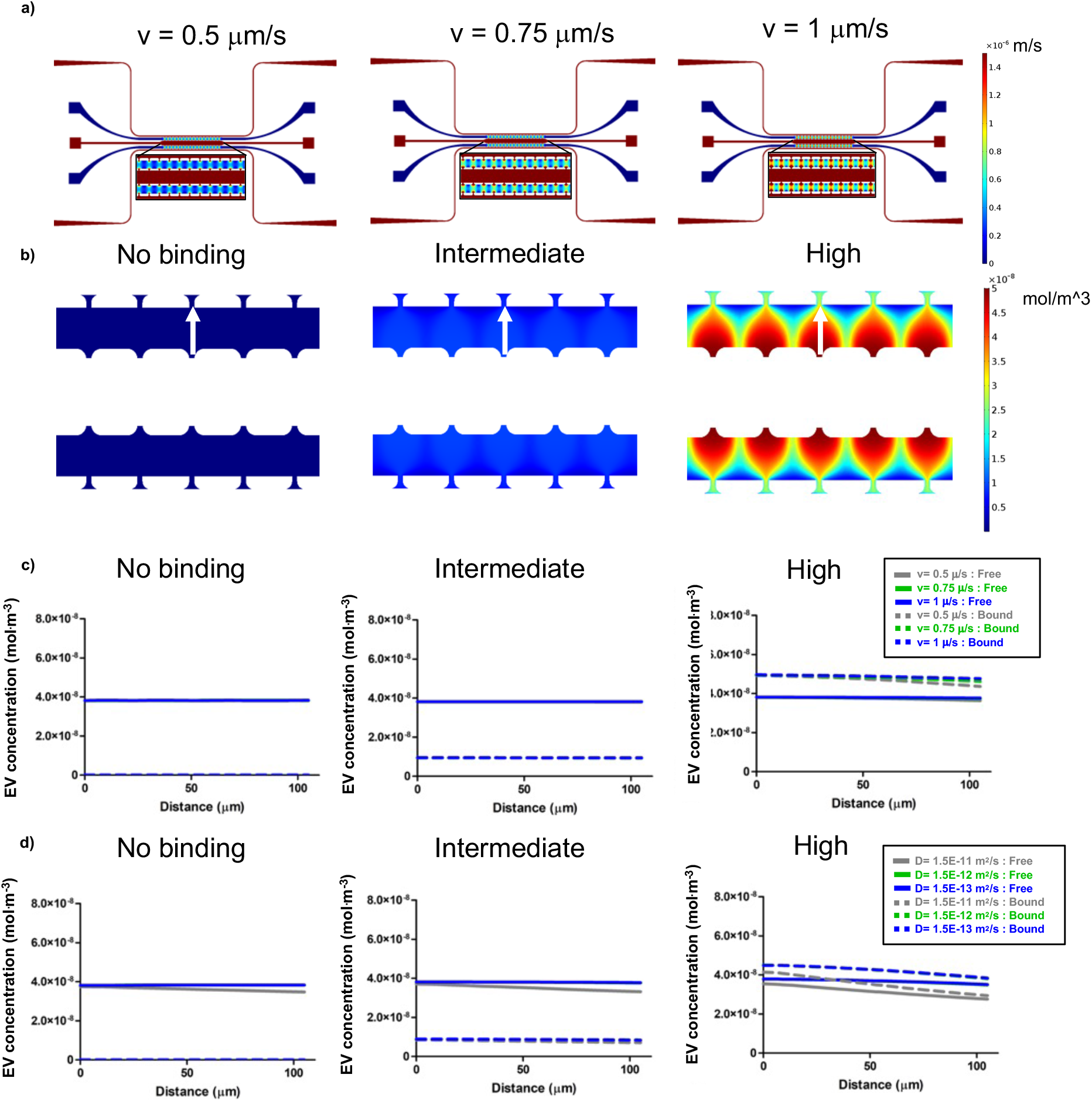
Finite element models reveal interstitial accumulation of high affinity binding EVs. a) A range of physiologically relevant interstitial flow velocities (v, 0.5 – 1 mm/s) were established by varying hydrostatic pressure boundary conditions at port inlets and outlets. b) Bound EV concentration profiles after T=30 min with a 0.5 mm/s flow velocity reveal accumulation of bound EVs as binding affinity increases. Concentration line profiles (white arrows) demonstrated c) flatter spatial EV concentration profiles with increasing flow velocity as well as higher levels of bound EVs with high affinity binding parameters. d) Comparing diffusivity of a small molecule dextran with EVs demonstrated differences in spatial profiles, but no significant difference when D < 10^-12^ m^2^/s which signifies EV transport is convection-limited.

We next examined the impact of molecular diffusion by simulating the convective transport into the interstitial channel using a range of molecular diffusivities that included a small molecular weight (~ 40 kDa) dextran (*D*=1.5 x 10^-11^ m^2^/s) in a hydrogel, approximate FRAP values of EVs in the laminin-rich ECM (*D*=1.5 x 10^-12^ m^2^/s), and an approximate value from an earlier report ^20^ that used particle tracking in a different hydrogel (*D*=1.5 x 10^-13^ m^2^/s). It is clear from all conditions, particularly those that include binding to the ECM, that diffusion of EVs (*D*=10^-12^ m^2^/s and smaller) does not impact the bound or free concentrations (1.5 x 10^-12^ and 1.5 x 10^-13^ m^2^/s curves completely overlap) (**Fig. 3d**). This result can be better understood by examining the relative rates of convection to diffusion using the dimensionless Peclet number (Pe), defined as *lv/D*, where *l* is a characteristic length of diffusion (100 μm channel and also similar to the distance between capillaries *in vivo*), and *v* is interstitial velocity. For a small dextran (~ 40 kDa), Pe= ~1.4 consistent with both convection and diffusion impacting net transport, which is evident in the shift in the spatial concentration profile when *D* is further decreased under all binding conditions (**Fig. 3d**). In contrast, for a typical EV (diameter of ~ 100 nm, *D*=1.5 x 10^-12^ m^2^/s) under the same conditions, Pe= ~ 33, and thus transport is dominated by convection (diffusion is negligible). This is also evident by the observation that any further decrease in *D* does not impact the bound or free concentrations of EV (**Fig. 3d**).

Finally, we explored the transient nature of the EV free and bound concentration by fixing *D* and v at our measured and physiologic values of 1.5 x 10^-12^ m^2^/s and 0.5 μm/s, respectively, and examined the spatial concentration profiles at different time points (0-60 min). Spatial gradients of both free and bound EVs are significantly attenuated over 60 min (**Fig. S5**). This observation suggests that steady state spatial concentration profiles are likely flat, and that binding of the EV to the ECM has a significant impact on the absolute concentration of EV. This conclusion is strongly dependent on our boundary condition at the lower pressure side of the interstitial chamber which assumes that all of the EVs are rapidly absorbed (e.g., by a lymphatic or blood vessel).

### *In vitro* EV convective transport and binding

We next sought to experimentally observe the transport of EVs under physiologically-relevant conditions consistent with our earlier experimental observations of diffusion and binding and *in silico* predictions (i.e., interstitial flow and a laminin-rich ECM). We utilized the same microfluidic device and conditions described for our *in silico* experiments (**Fig. S4**), and the same ECM and populations of EVs (from the MCF10 series) and experimental conditions from the diffusion-binding experiments.

To validate experimental interstitial flow velocity (i.e., flow through chambers 2a and 2b, **Fig. S4**)), fluorescently labeled 40kDa dextran was added to PBS in chamber 1. Laminin-rich ECM polymerization is anisotropic ^45^ resulting in spatially heterogeneous hydraulic conductivity and thus interstitial flow. To account for this, we measured the convective transport of the non-binding dextran at each port (**Fig. S6a**), and then used our *in silico* model to estimate the local permeability and thus interstitial velocity at each pore (**Fig. S6b,c**). The interstitial velocity at most microfluidic device pores fell in a range (0.15-0.75 μm/s; mean = 0.29 μm/s) that was physiologic and consistent with our *in silico* experiments. Pores outside of this range were not included in the subsequent analysis.

Cell line (i.e., malignant potential) and integrin binding state significantly impacted the concentration and spatial distribution of EVs at time 30 minutes (**Fig. 4, supplementary videos**). Accumulation was most apparent within the first 100 μm of the laminin-rich ECM (**Fig. 4b**; white arrows). Highest EV accumulation was observed with MCF10CA1 EVs treated with MnCl_2_ (**Fig. 4b**; right column), which is consistent with high binding observed in FRAP and *in silico*. The addition of EDTA or integrin blocking antibodies (**Fig. 4c,d**) neutralized EV accumulation, particularly with the malignant MCF10CA1 EVs, but also for normal MCF10A and partially malignant MCF10DCIS EVs.

**Figure 4.**
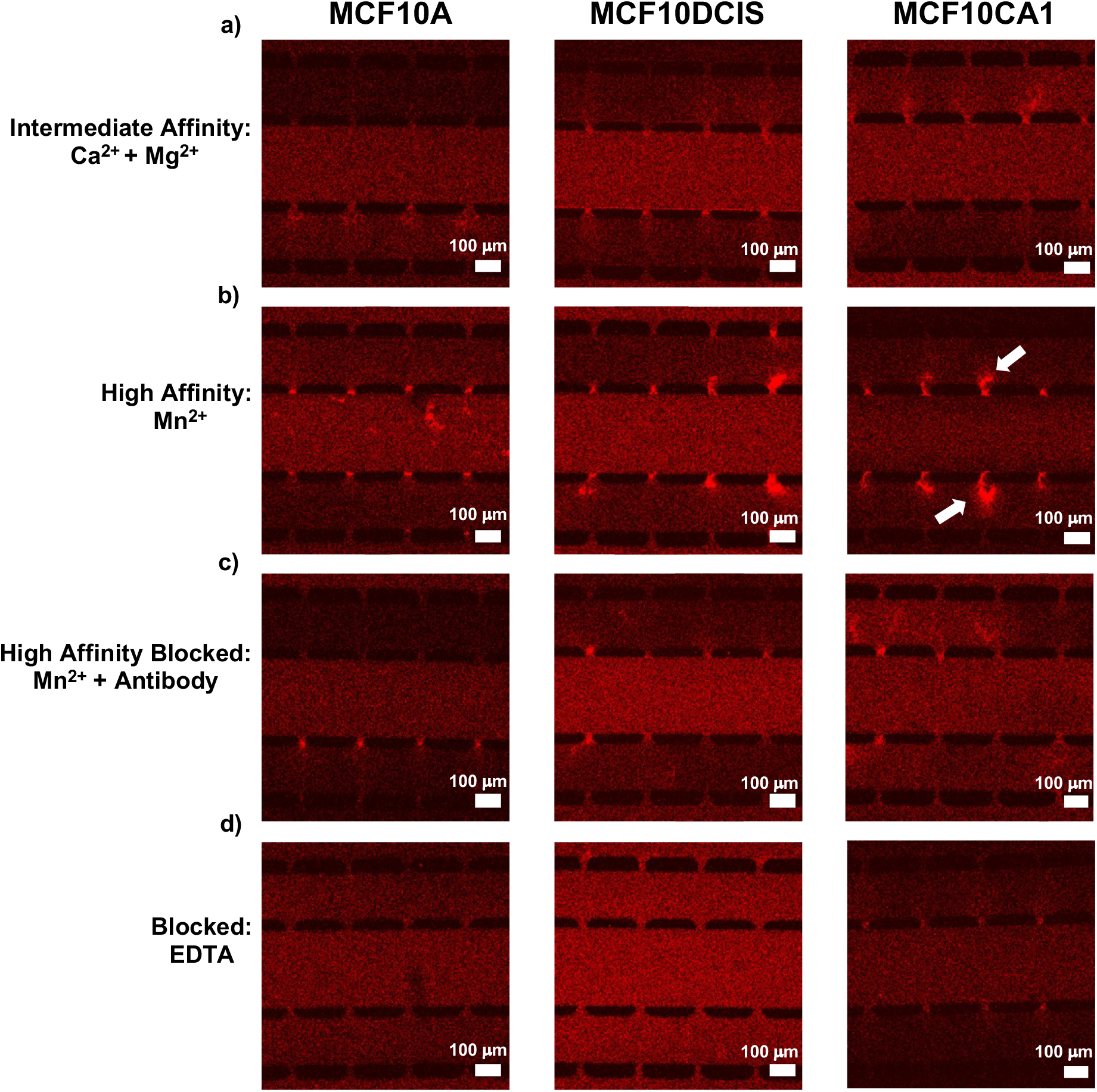
Interstitial EV accumulation with increased malignancy and high affinity integrin binding. **a-d)** Representative images demonstrated a range of EV binding profiles. Most consistent high binding was observed with MCF10CA1 EVs treated with Mn2+ (b, right column, white arrows) for high affinity binding. Differences in background fluorescent intensity were evident but were normalized for quantitative analysis.

To quantify differences between cell lines and experimental conditions, the fluorescent intensity in the interstitial chamber (chamber 2), after 30 minutes of convective flow, was first normalized to that in the central chamber (the source, chamber 1). The normalized fluorescent intensity was then determined along a linear region across the interstitial chamber (**Fig. 5a**), and resulting curves were averaged (**Fig. 5b-h**). High affinity MCF10CA1 EVs accumulated to 8-fold higher than source concentration followed by MCF10DCIS and MCF10A (**Fig. 5c**). Blocking conditions including an antibody cocktail against laminin binding integrins (**Fig. 5d)**, individual integrin blocking antibodies (**Fig. 5f,g**), or EDTA (**Fig. 5e**) dramatically reduced EV accumulation of MCF10CA1 and MCF10DCIS EVs and to a lesser extent MCF10A EVs. Intermediate affinity conditions demonstrated similarly low levels of EV accumulation across cell lines (**Fig. 5b,h).** These results demonstrate that EV binding affinity directly impacts the interstitial spatial distribution and absolute concentration, and that the binding is partially dependent on integrins α3, α6, and β1.

**Figure 5.**
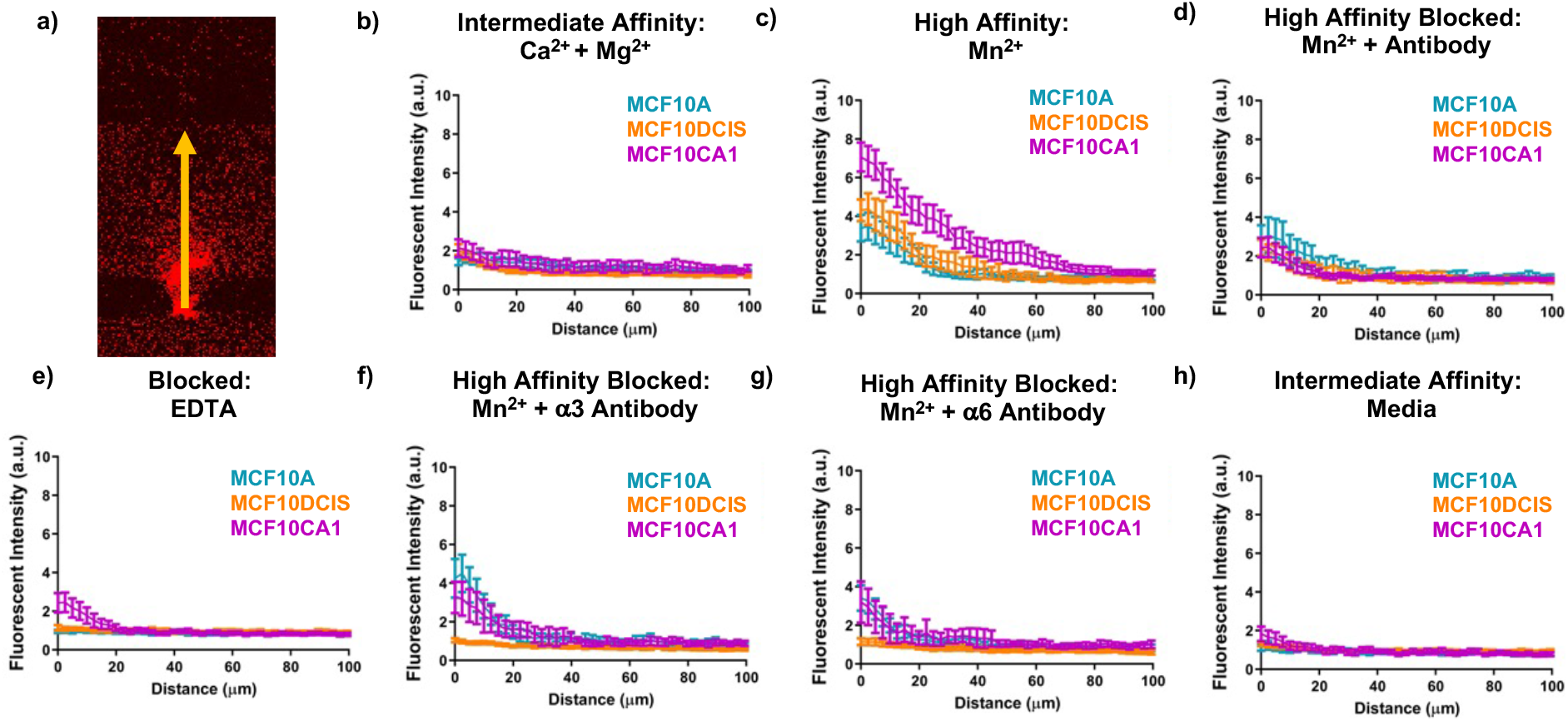
EV convective transport is affected by parent cell malignancy and integrin activation state. a) Line profiles were used to assess EV concentration after T=30 min of flow. b-h) Comparisons of concentration profiles by integrin activation state showed highest differences for high affinity MCF10CA1 and MCF10DCIS EV binding. Blocking conditions reduced the formation of spatial gradients. Physiologic binding conditions showed minimal differences between EV conditions. Error bars=95% CI. n=2-4 devices per curve (average devices per condition=3); n=21-80 ports per curve (average ports per curve=52).

## Conclusions

EVs can impact biology in myriad ways including critically important phenomena such as cancer metastasis. In addition, the inherent qualities of EVs provide an exciting and flexible new platform for novel nanotherapeutics. Progress has been slowed by a limited understanding of the fundamental mechanisms of transport in tissue spaces, which will dictate the spatial distribution and concentration. Using a combination of technologies, we demonstrate that diffusion of EVs under physiological conditions in the interstitium is negligible, and that transport, and thus the spatial distribution of bound and free EVs, is dominated by interstitial flow (convection) and binding to the extracellular matrix. The EVs, derived from a cell line series of progressively increasing malignancy, express laminin-binding integrins (α3, α6, and β1) which contribute to the spatial distribution and concentration of EVs. These results provide a paradigm shift in our understanding of the mechanisms that control interstitial transport of EVs and should impact the design and strategies to deliver nanotherapeutics broadly.

## Acknowledgements

This work supported, in part, by an NSF Graduate Research Fellowship (PAS), the ARCS Foundation Scholar Award (PAS), grants from the National Institutes of Health (UH3 HL141800, R01 EB030410, R21 AI161041, R01 CA241666, F31 NS120590), and funds from the Department of Biomedical Engineering and the College of Engineering at the University of California, Davis.

## Contributions

P.A.S, R.R.M, V.S.S., D.M.R., R.P.C. and S.C.G. conceptualized and planned the experiments, and participated in the data analysis and interpretation. P.A.S, R.R.M, A.B., C.N., and B.S. performed the experiments. P.A.S. and S.C.G. prepared the figures. P.A.S, V.S.S., R.R.M., A.B., C.N., B.S., D.M.R., R.P.C. and S.C.G. wrote the manuscript, and all authors reviewed and edited the manuscript before submission.

## Ethics declarations

There are no conflicts of interest to declare.

## Materials and Methods

### Cell culture

The MCF10 breast cancer progression series (MCF10A, normal; MCF10DCIS, pre-malignant; MCF10CA1, malignant) was purchased from Karmanos Cancer Center, and sub-cultured according to advised protocols. Culture medium for MCF10A cells consisted of DMEM/F12 (Thermo Fisher Scientific), supplemented with 5% horse serum (Thermo Fisher Scientific), 1.05 mM CaCl_2_ (Sigma-Aldrich), 20 ng/mL EGF (Thermo Fisher Scientific), 105 ng/mL cholera toxin (Millipore Sigma), 10 μg/mL insulin (Sigma), 0.5 μg/mL hydrocortisone (StemCell Technology). MCF10DCIS and MCF10CA1 culture medium consisted of DMEM/F12 supplemented with 5% horse serum and 1.05 mM CaCl_2_.

### EV isolation

To prepare EV-depleted serum for culture media, serum-depleted horse serum was prepared by serial centrifugation. Serum was centrifuged for 300xg for 10 minutes at 4°C to remove live cells and large debris, 2,000xg for 15 minutes at 4°C to remove dead cells and apoptotic bodies, 10,000xg for 30 minutes at 4°C to remove larger microvesicles, and finally spun via ultracentrifugation overnight at 120,000xg at 4°C. Absence of serum EVs was validated via nanoparticle tracking analysis (NTA) with the NanoSight LM10 (Malvern Panalytical Ltd.).

To isolate MCF10 series EVs, MCF10A, MCF10DCIS, and MCF10CA1 cells were loaded into 3 T-150 flasks each at a starting confluency of approximately 15% (1.5×10^6^ MCF10A cells, 1.5×10^6^ MCF10DCIS cells, 2×10^6^ MCF10CA1 cells) and cultured for 24 hours in non-EV depleted media. Following 24-hour culture, flasks were triple rinsed with ample PBS + 1mM CaCl_2_ and 1 mM MgCl_2_ (Thermo Fisher Scientific) to remove horse serum-derived EVs. 15 mL of EV-depleted serum media was added to each flask and cells were cultured for 72 hours. Final cell concentration was observed following 72-hour culture and was between 80-95% confluent. Initial seeding was optimized to prevent 100% confluence over the course of the 4-day culture.

Conditioned media was collected and subjected to increasing centrifugation spins (300xg for 10 minutes at 4°C, 2,000xg for 15 minutes at 4°C, 10,000xg for 30 minutes at 4°C). 150,000 kDa Amicon filters (Millipore Sigma) were loaded with 15 mL of pre-spun conditioned media and spun at 4,000xg for 45 minutes according to manufacturer protocols for crude EV purification. The EV-retentate was collected and filters were washed with MilliQ water. EVs for downstream imaging analysis were stained with 2 mM CellTrace Far Red (Thermo Fisher Scientific) and incubated for 2 hours at 37°C ^46^.

Following incubation with CellTrace Far Red (CTFR), EVs were separated from excess dye as well as free proteins via size exclusion chromatography (SEC). An Izon Automatic Fraction Collector and qEV_original_ 35 nm SEC columns (Izon) were loaded with 1 mL of concentrated EV stock. EV-rich fractions one through four were collected (0.5 mL each) and utilized for downstream experiments, while fractions five through twelve were collected to serve as EV-free controls.

### Nanoparticle tracking analysis (NTA)

EV samples were diluted 1:250 in 0.22 μm filtered MilliQ water, and 1 mL samples were added to a NanoSight LM10 (Malvern Panalytical Ltd) to characterize EV size and concentration. An automated syringe pump (Harvard Bioscience) provided consistent flow of EV sample across the field of view to provide multiple measurements. 3 x 30s videos were acquired per sample at camera level 13, and NanoSight NTA 3.1 software was utilized for analysis with detection threshold 3-4.

### Transmission electron microscopy (TEM)

EVs were fixed in 1% glutaraldehyde for 5 minutes. A 10 μL droplet of EV/ glutaraldehyde mix was placed on parafilm and a copper formvar grid was floated, copper side down, on the droplet for 40 minutes. The grid was then dried on filter paper, moved to a 100 μL droplet of MilliQ water to wash, dried again, and moved to a 50 μL droplet of 0.2 um filtered 4% uranyl acetate for 8 minutes. Grids were then dried again on filter paper and allowed to air dry for at least 10 minutes before imaging. Grids were imaged on a Talos FEI L120C TEM (Thermo Fisher Scientific) from 11,000x to 36,000x magnification.

### Single EV protein characterization-ExoView

An ExoView R100 (NanoView Biosciences) was utilized for individual EV immunofluorescent analysis. EVs were diluted to an initial concentration between 6.66E6-1.32E7 EV/mL and diluted 1:1 with incubation solution (NanoView Biosciences) according to manufacturer protocols. EV solutions were loaded onto EV-TETRA-C ExoView Kits (NanoView Biosciences) and prepared via an ExoView CW100 Chip Washer (NanoView Biosciences). Fluorescently conjugated integrin antibodies (1:400 AF488 anti-integrin β1, clone-TS2/16, BioLegend; 1:200 PE anti-integrin β1, clone-TS2/16, BioLegend; 1:400 AF647 anti-integrin β1, clone-TS2/16, BioLegend; 1:400 PE anti-integrin α3, clone-ASC-1, BioLegend; 1:400 AF488 anti-integrin α6, clone-GoH3; BioLegend) and ExoView tetraspanin kit antibodies (1:500 AF647 anti-CD63, clone-H5C6; 1:500 AF555 anti-CD81, clone-JS 81; 1:500 AF488 anti-CD9, clone-HI9a) were added at the appropriate steps during the incubation procedure. MIgG thresholds were selected at approximately the 75^th^ percentile to reject the majority of non-specific binding. EV counts and integrin or tetraspanin colocalization was normalized to MigG counts.

### Microfluidic platform

The microfluidic device leveraged for diffusion, convection, and *in silico* experiments was previously developed in the lab ^23^. This device consists of five parallel chambers communicating with other by diffusion and convection through small evenly spaced pores (**Fig. S4**). Microfluidic devices were fabricated with via soft-lithography with polydimethylsiloxane (PDMS). PDMS which was prepared by mixing a 10:1 ratio of Sylgard™ 184 silicone elastomer base and curing agent (Dow Corning) and pouring over SU-8 master molds. Cast-PDMS devices were bonded to glass coverslips (Thermo Fisher Scientific) via plasma treatment (Harrick Plasma).

200 μL pipette tips (Genesee Scientific) were added to fluidic ports at the end of chambers 1, 3a, and 3b, and filled with varying fluid volumes to establish controlled hydrostatic pressure heads. Differences in hydrostatic pressure head heights were leveraged to drive convective flow in *in silico* and *in vitro* convective flow experiments. Communication between chambers was established by structural micropores, which were previously optimized to avoid leakage of gels into neighboring chambers while also providing air-bubble free gel/fluid interfaces (pore width=30 μm; pore length=550 μm).

### *In vitro* diffusion experiments

To assess diffusion, CTFR stained MCF10 series EVs (1E9 EVs) were incubated with 1 mg/mL BSA (Thermo Fisher Scientific) for 15 minutes at 4°C to block nonspecific binding interactions. EVs were then incubated with a working solution of either 0.1 mM CaCl_2_ and 1 mM MgCl_2_, 1 mM MnCl_2_, 0.1 mM CaCl_2_ and 1 mM MgCl_2_ (all from Sigma-Aldrich) with a cocktail of functionally inhibitory integrin blocking antibodies (20 μg/mL anti-α3, clone-ASC-6, Millipore Sigma; 20 μg/mL anti-α6, clone-GoH3, Thermo Fisher Scientific; 50 μg/mL anti-β1, clone-mAb13, Millipore Sigma; 20 μg/mL anti-β4, clone-ASC-6, Millipore Sigma), or 1 mM MnCl_2_ with the cocktail of functionally inhibitory integrin blocking antibodies. EV solutions were then mixed with laminin-rich ECM (Matrigel-GF Reduced; Corning) for a final laminin-rich ECM working concentration of 3 mg/mL and loaded in chamber 2a or 2b (**Fig. S4**). This concentration of Matrigel results in a matrix with a range of pore sizes similar to that present *in vivo* (50-5000 nm) ^40^. Laminin-rich ECM was left to polymerize for 30 min at 37°C before FRAP imaging. Care was taken to prevent bulk convective fluid flow by ensuring loading heads were roughly equivalent, incubating for an additional 15 minutes at room temperature to allow time for fluid height equilibration, and leaving device chambers 1, 3a, and 3b devoid of fluid.

### Fluorescence recovery after photobleaching (FRAP)

FRAP was performed on an Olympus FV3000 laser scanning confocal microscope (Olympus) via a 20x objective with 3x digital zoom and captured by high sensitivity-spectral detector. FRAP images were acquired every 240 ms over the course of 5 minutes with a 100% laser excitation pulse after the first four frames to generate a circular 18 μm diameter bleach spot. Laser power was tuned to ensure that the final non-bleached background fluorescence was within 15% of the initial non-bleached background fluorescence on average.

The underlying pseudo first order kinetic equation describing EV interactions with the ECM can be described by:

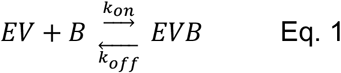

where EV is the concentration of free EVs, B is the concentration of binding sites within the matrix, and EVB is the concentration of matrix-bound EVs. Initial concentrations of each species as well as forward (k_on_) and reverse (k_off_) rate constants determine final steady state concentrations of each species. The ratio of k_off_/kon provides the dissociation constant, K_d_.

Raw FRAP data was analyzed in FIJI using a standard analysis pipeline. In brief, the “create profile” plugin ^47^ was implemented to generate average fluorescent intensity spectrum curves over the full image stack. ROIs were generated for the bleach spot as well as a control ROI for the background fluorescence. The bleach spot fluorescent spectrum was normalized to a function fit to the background fluorescence ROI to control for photobleaching. Spectrum data was exported to MATLAB for further analysis. To perform fits, background-normalized spectrum data was normalized to fall within 0 and 1 by setting the mean value pre-bleach equal to 1 and the minimum value post-bleach equal to 0. A progressive sliding window was utilized to average timepoints in order to smooth the curves and to prevent biased overfitting of later timepoints. Sliding window conditions were as follows: T=0-0.72s (Pre-bleach) - no averaging; T=0.96-29.76s – averaged 2 points; T=30.76-44.4s – averaged 5 points; T=46.8-58.8s – averaged 10 points; T=62.4-66s – averaged 15 points; T=70.8-299.76s – averaged 20 points. Log space parameter arrays with 50 values were generated to screen the three fitting parameters: D_eff_ (1E-13->1E-11), K*on (1E-7->1E-2), and K_d_ (1E-6->1E-1).

FRAP curve fitting procedures followed the general methodology previously described in detail to fit experimental data with pseudo first order kinetics (Eq. 1). The following fitting equation was applied to the experimental FRAP recovery curves ^39^:

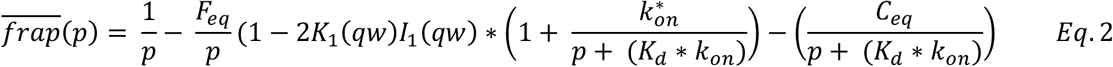

where,

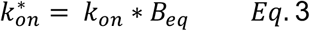

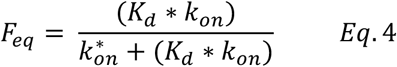

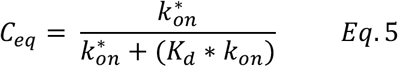

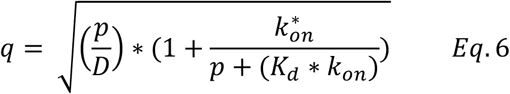

where 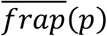 is the average of the Laplace transform of the fluorescent intensity within the bleach spot, p is the Laplace variable that inverts to yield time, I_1_ and K_1_ are modified Bessel functions of the first and second kind respectively, w is the bleach spot radius, B_eq_ is the equilibrium concentration of binding sites, and D is the EV free diffusion coefficient. The numerical inversion of this function was performed via the “invlap.m” routine.

Fitting for exact values of k*_on_ and k_off_ from fluorescence recovery curves is difficult due to the long relatively flat trough observed when plotting the minimized sum of squares of the residual (SSR) as a function of k*_on_ and k_off_. In other words, multiple combinations of k*_on_ and k_off_ produce similar SSR, thus rendering it impossible to uniquely identify the individual binding parameters. The solution is to simply fit for the ratio of two, or K_d_ (K_d_=koff/kon) (**Fig. S3**). A matrix of 50×50×50 curve libraries was generated by importing each combination of D, k*_on_, and K_d_ fitting parameters. A sum of squares approach was applied to minimize the error between library curves and experimental data. Preliminary best-fit parameters were extracted from the best fit curve.

To refine the fitting procedure, the MATLAB routine “fitnlm”, was applied for higher accuracy fitting of k*_on_ and K_d_. D was constrained to the value determined through the first sum of squares residual estimation to reduce the burden on the fitting process and risk of overfitting the data. The “fitnlm” routine used the Levenberg-Marquardt nonlinear least squares algorithm. Resulting refined-fit parameters for k_on_ (k*_on_/B_eq_) and K_d_ are presented in results, although the difference between the initial sum of squares fit and the refined fit were negligible.

### Finite element simulations (*in silico* experiments)

Computational finite element simulations were performed using COMSOL Multiphysics™ 5.2a software. The convection-diffusion equations of mass and momentum transport were solved to find flow velocities and spatial concentration profiles of EVs, binding sites, and bound EVs. The Free and Porous Media Flow module was used to model fluid through the porous ECM, and Transport of Diluted Species Modules were implemented to track transport of each species. Chemistry modules were added to capture the relationship between EVs and binding sites (Eq. 1). 2D solutions were generated assuming incompressible, single-phase, laminar flow with no-slip boundary conditions applied to all surfaces with the exception of terminal port inlets and outlets (ends of chambers 1-3). EVs were modeled as a dilute, dissolved species which neglected physiologic drag and interactions with matrix porosity. Binding sites and bound EVs were modeled as effectively immobile with *D* = 1E-16 m^2^/s. Baseline modeling parameters (**Suppl. Table 2**) were used unless otherwise specified. Additional parameter values were: dynamic viscosity of water at 25°C = 0.89 cP; density of water at 25°C = 1000 kg/m^3^; porosity of matrix = 0.99 %. Various parameter and time sweeps were performed, and parameter values were tested iteratively along with *in vitro* experiments.

### *In vitro* interstitial flow experiments

Identical microfluidic devices as those used in FRAP experiments were utilized for convective flow experiments. CTFR-stained EVs were prepared as described above but were not embedded in laminin-rich ECM. Rather, laminin-rich ECM was prepared at 3 mg/mL and loaded into chambers 2a and 2b. Following a 30-minute incubation at 37°C, warmed PBS was added to chambers 1, 3a, and 3b to establish hydrostatic pressure heads and convective flow from chamber 1 outwards across the matrix-filled chambers 2a and 2b towards fluidic chambers 3a and 3b. To initiate fluid flow for convective flow experiments the following fluid volumes were added to 200μl pipette tips to establish hydrostatic pressure heads: Chamber 1: Inlet-35μl, Outlet-30μl; Chamber 2a,b: filled with laminin-rich ECM, no additional pressure head was applied ; Chamber 3a,b: Inlet-15μl, Outlet-10μl. Pressure heads were selected to ensure symmetry from the middle of chamber 1 outwards to fluidic chambers 3a and 3b.

40 kDa FTIC dextran (Sigma-Aldrich) was added to EV solutions to serve as a control for fluid flow. EV and dextran flows were visualized via an Olympus FV3000 confocal microscope (Olympus) with a 10x objective using 647nm and 488nm lasers and captured by high sensitivity-spectral detectors to maximize capture of the EV fluorescent signal. Dextran velocity was calculated using iterative *in vitro* experiments and finite element modeling. In brief, displacements of dextran fluorescence over time were compared between *in vitro* and computational models. *In silico* parameter sweeps of ECM permeability provided associated interstitial flow velocity values. Curves were generated to plot ECM permeability vs. dextran displacement and interstitial fluid velocity vs. ECM permeability. *In vitro* dextran displacement was plotted along these curves to identify interstitial flow velocity.

To assess EV convective flow and accumulation in the matrix, the following solutions were added to each device chamber via the same methodology and volumes indicated above: Chamber 1: EV + 40 kDa dextran + treatment condition + PBS ; Chamber 2a,b: filled with laminin-rich ECM ; Chamber 3a,b: PBS. EV treatment conditions used the same concentrations of blocking antibodies described in diffusion experiments. 5mM EDTA (Thermo Fisher Scientific) and RPMI 1640 were added for additional treatment conditions.

Images were acquired at 1.08 second intervals over the first 10 minutes of flow. After 30 minutes of flow, images of the full device were acquired as a terminal endpoint. Data for convective flow experiments was analyzed in FIJI. Fluorescent intensity line profiles were generated to show EV concentration gradients across laminin-rich ECM chambers. Multiple ports from each device were measured, but analysis was limited to ports with flow velocities within a binned range: 0.15 μm/sec ≤ v ≤ 0.75μm/sec. Binning was performed to exclude ports with negligible flow as well as those with rapid flow caused by degraded laminin-rich ECM. The majority of ports were within the binned range.

### Statistical analysis and figure generation

The majority of experimental conditions were performed in triplicate at minimum. PRISM 9.2 was utilized for statistical analysis and generation of graphs. One-way ANOVA tests with a p-value < 0.05 and Tukey-Post hoc tests were employed when comparing three or more conditions, while student T-tests with a p-value < 0.05 were used to show significance when comparing only two conditions.

## Tables

**Supplementary Table 1.** The percentage of MCF10 series EVs with colocalized integrins.

|  | MCF10A | MCF10DCIS | MCF10CA1 |
| --- | --- | --- | --- |
| CD63 Capture |  |  |  |
| $\alpha 3\beta 1$ | 7% | 23% | 24% |
| $\alpha 6\beta 1$ | 5% | 6% | 20% |
| CD9 Capture |  |  |  |
| $\alpha 3\beta 1$ | 19% | 27% | 45% |
| $\alpha 6\beta 1$ | 8% | 11% | 39% |

**Supplementary Table 2.** Baseline modeling parameters used for *in silico* finite element simulations.

| Parameter | Value | Units | Source |
| --- | --- | --- | --- |
| Diffusivity (D) | 1.53E-12 | m<sup>2</sup>/s | -Approximated from experimental FRAP results and literature <sup>20</sup> |
| Concentration of binding sites | 4.2E-3 | mol/m<sup>3</sup> | -Calculated from concentration of 3 mg/mL laminin-rich ECM with one binding site per laminin monomer <sup>48</sup> |
| EV influx concentration | 4.16E-8 | mol/m<sup>3</sup> | -Experimentally determined as the minimum observable concentration on the confocal microscope |
| Interstitial flow velocity | 0.5E-6 | m/s | -Approximate general value for interstitial flow <sup>49,50</sup> |
| Permeability | 2E-14 | m<sup>2</sup> | -Experimentally determined for 3mg/mL laminin-rich ECM |
| Binding States | $k_{on}$ (m<sup>3</sup>·s/mol) | $k_{off}$ (1/s) | $K_d$ (mol/m<sup>3</sup>) |
| No binding (EDTA or antibody blocked) | 0 | 0 | n/a |
| Intermediate affinity (10A Ca<sup>2+</sup> + Mg<sup>2+</sup>) | 0.143 | 2.38E-3 | 16.6E-3 |
| High affinity (10CA1 Mn<sup>2+</sup>) | 0.542 | 1.65E-3 | 3.04E-3 |

## Supplementary Figures

**Figure S1.**
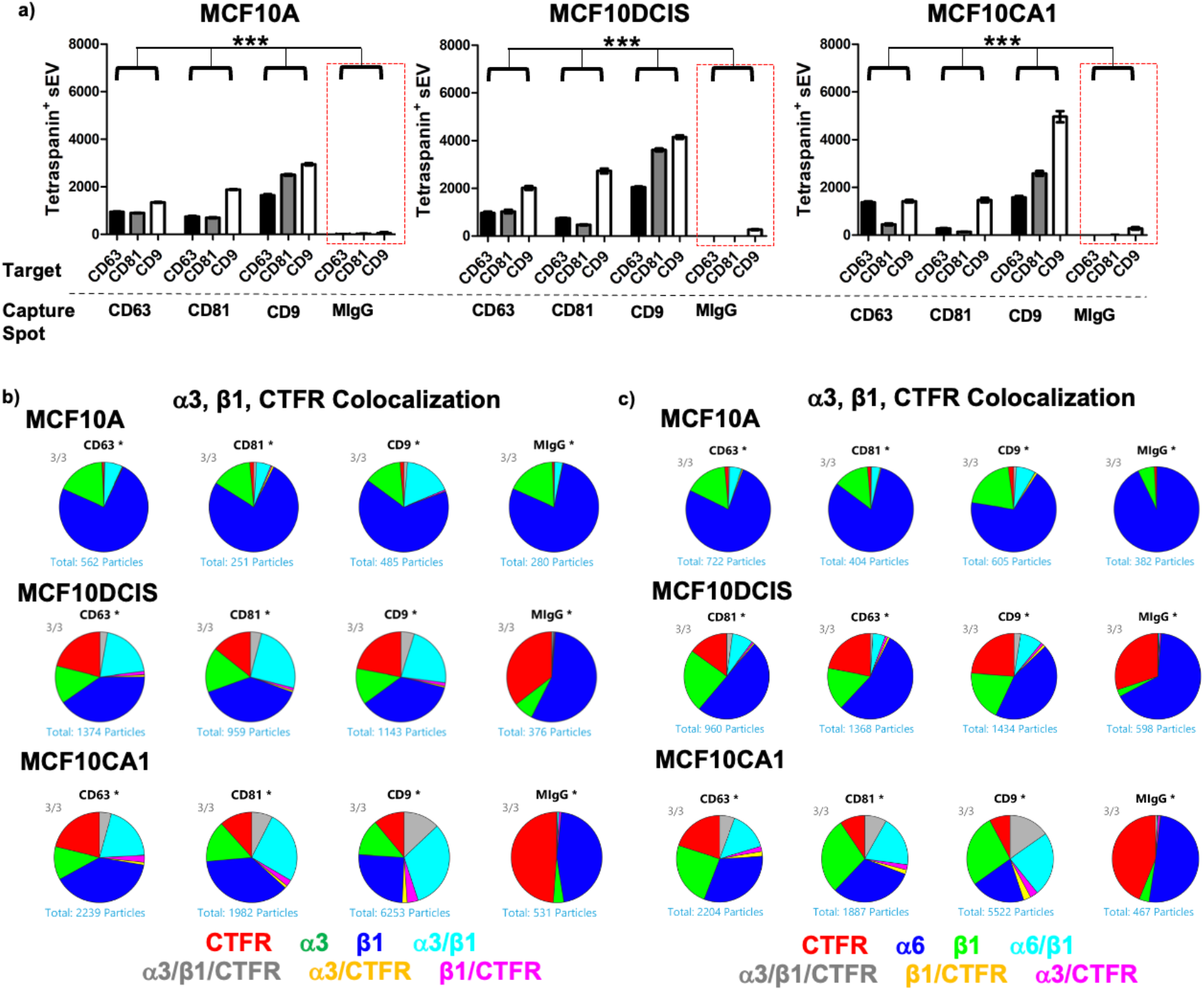
Isolated EVs possess appropriate tetraspanin markers and colocalized integrins. a) ExoView analyses revealed CD63, CD81, and CD9 tetraspanin presence on EVs isolated from each of the MCF10 series lines when compared to negative MlgG capture antibody control (red boxed). ***p<0.001; One-way Anova, Tukey post-hoc. b) Pie charts comparing CTFR^+^, integrin α3, and integrin β1 colocalization generally show higher percentages of colocalized markers on increasingly malignant EVs. c) CTFR^+^, integrin α6, and integrin β1 colocalization follows a similar trend. Although the same number of EVs (counted by NTA) were loaded, differences in captured EV counts were observed.

**Figure S2.**
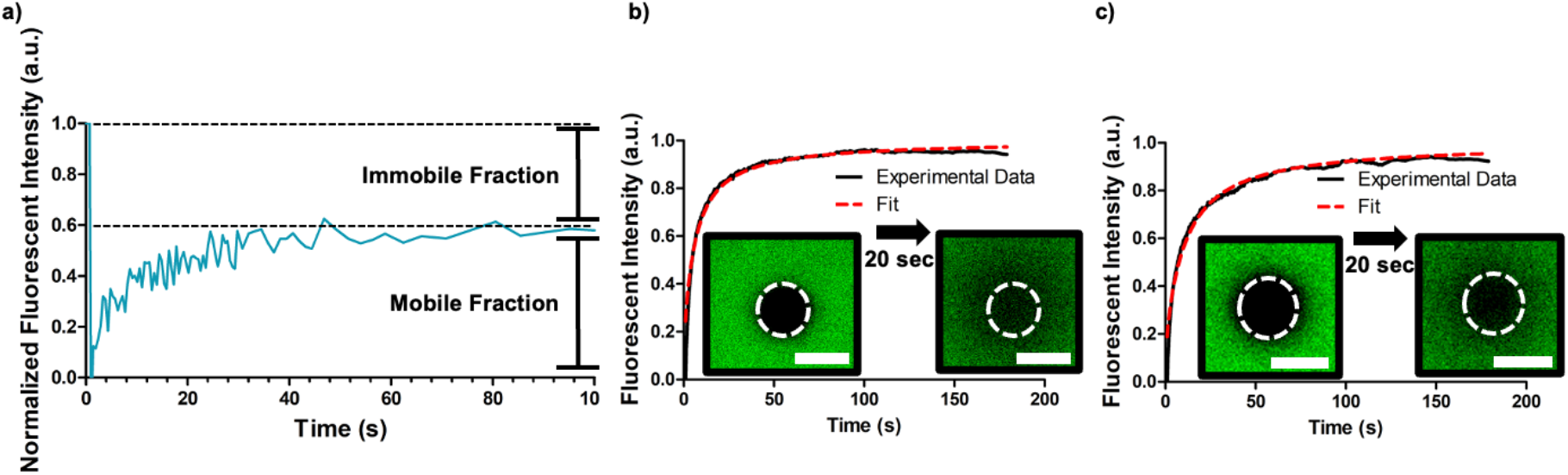
Characteristic fluorescence recovery after photobleaching (FRAP) curve. **a)** Fluorescent species (1 a.u.) are bleached with a high intensity laser (0 a.u.), and recovery dynamics are recorded in the bleached region of interest. Where the FRAP curve asymptotically recovers to in relation to pre-bleach intensity reveals the proportion of species that are freely mobile versus immobile. Fluorescence recovery of b) 40 kDa FITC dextran and c) 150 kDa FITC dextran, in 3 mg/ml laminin rich ECM. Representative bleach spot recoveries over 20 seconds. Scalebars=20 μm; n=6-9 averaged replicates per curve.

**Figure S3.**
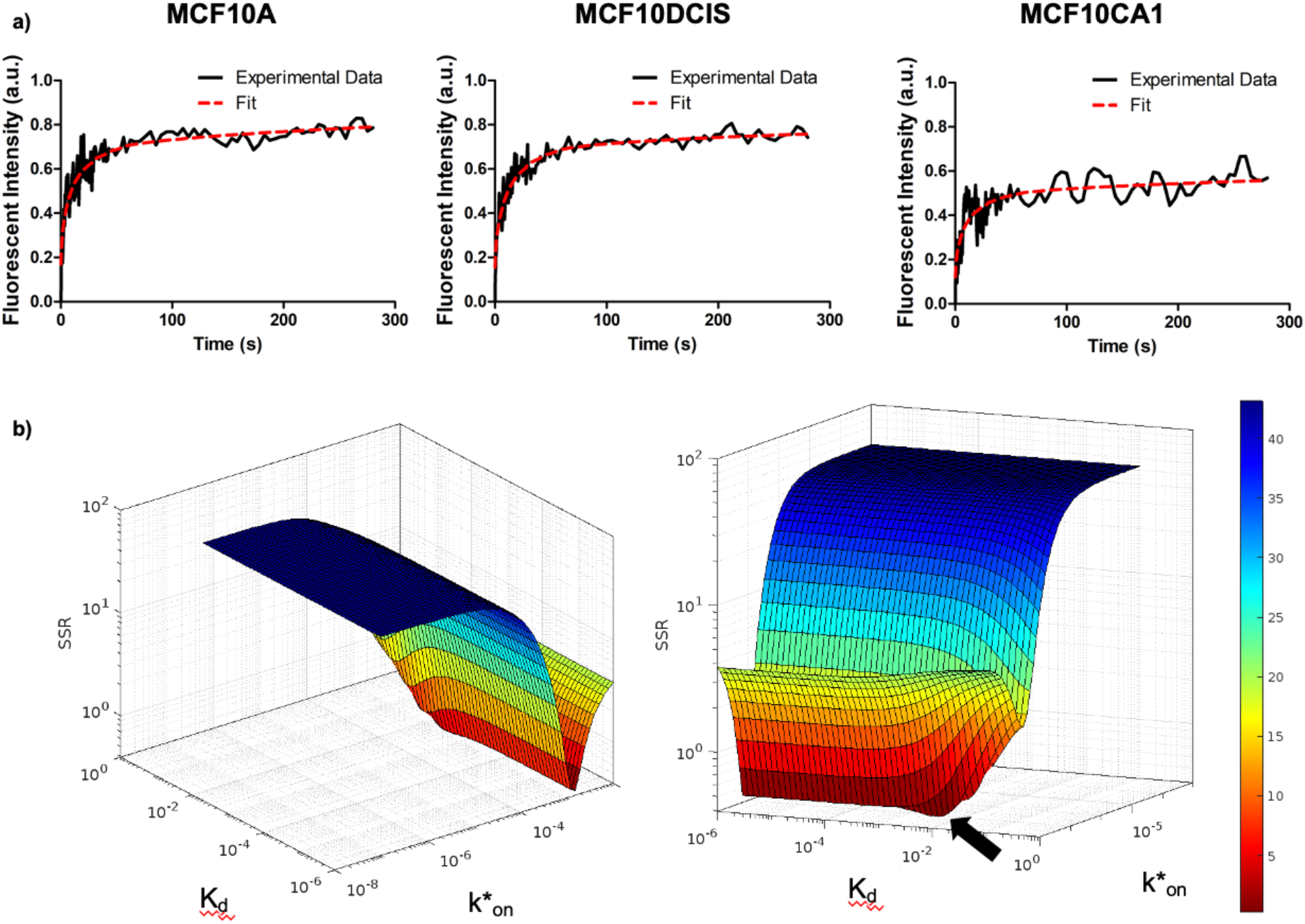
Fluorescence recovery after photobleaching (FRAP) curves of EVs can be fit to determine molecular diffusivity and equilibrium binding coefficients. **a)** Representative FRAP curves for EVs from each MCF10 line were fit to a diffusionreaction model, b) Surface plots displaying sum of squares residuals (SSR) as a function of the forward and dissociation rates (k*_on_ and K_d_) demonstrate a long relatively flat trough consistent with a range of combinations of k*_on_ and K_d_ which provide approximately the same fit. SSR minimum (black arrow) reflects the best fit k*_on_ and K_d_ pair.

**Figure S4.**
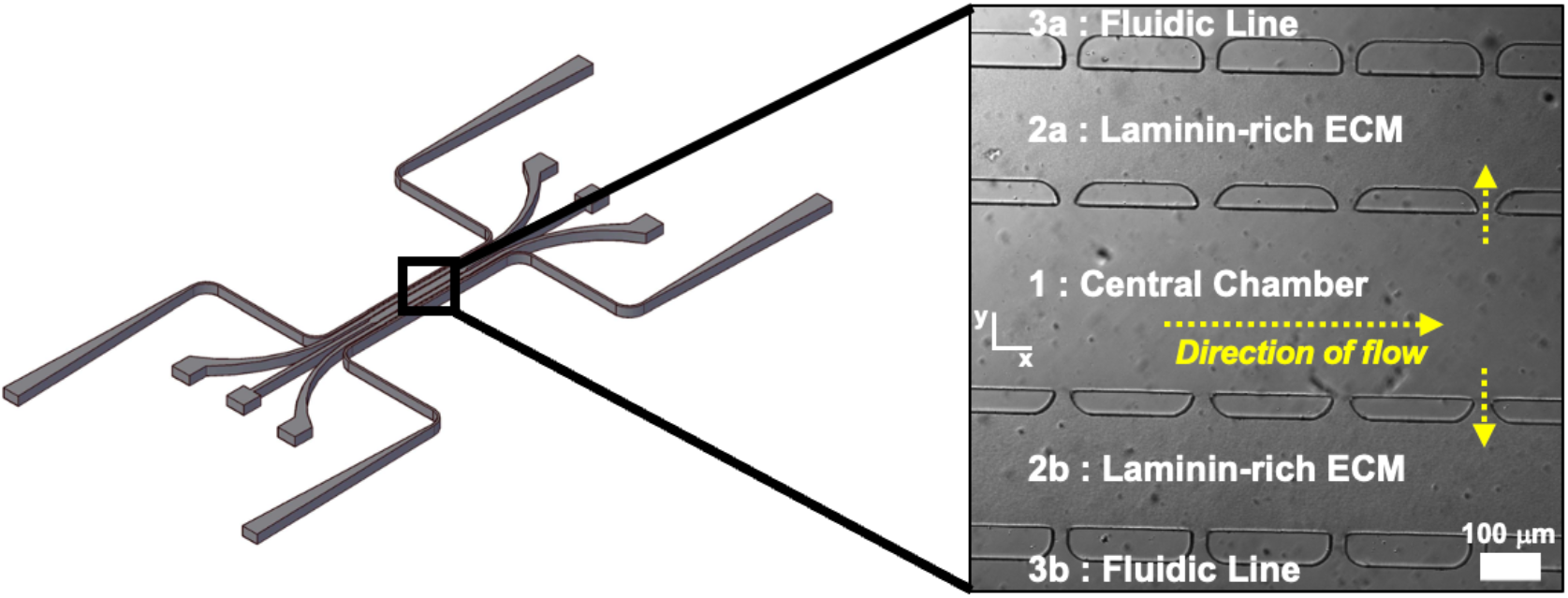
Microfluidic device for *in silico* and *in vitro* EV transport. The microfluidic platform features five parallel chambers that can communicate through small pores (diameter 30 microns). Pipette tips at the ends of each channel can establish hydrostatic pressures such that there are steady pressure gradients in the x and y directions. Compartments #2a and #2b are loaded with a laminin-rich ECM (3 mg/ml). The central chamber (#1) delivers a constant concentration of EVs by convection (left to right), and is at a higher pressure at each equivalent x-position than the adjacent compartments (#2a and #2b), which are at a higher pressure that the outer compartments (#3a and #3b). This creates a constant convective interstitial flow (y-direction) through compartment 2.

**Figure S5.**
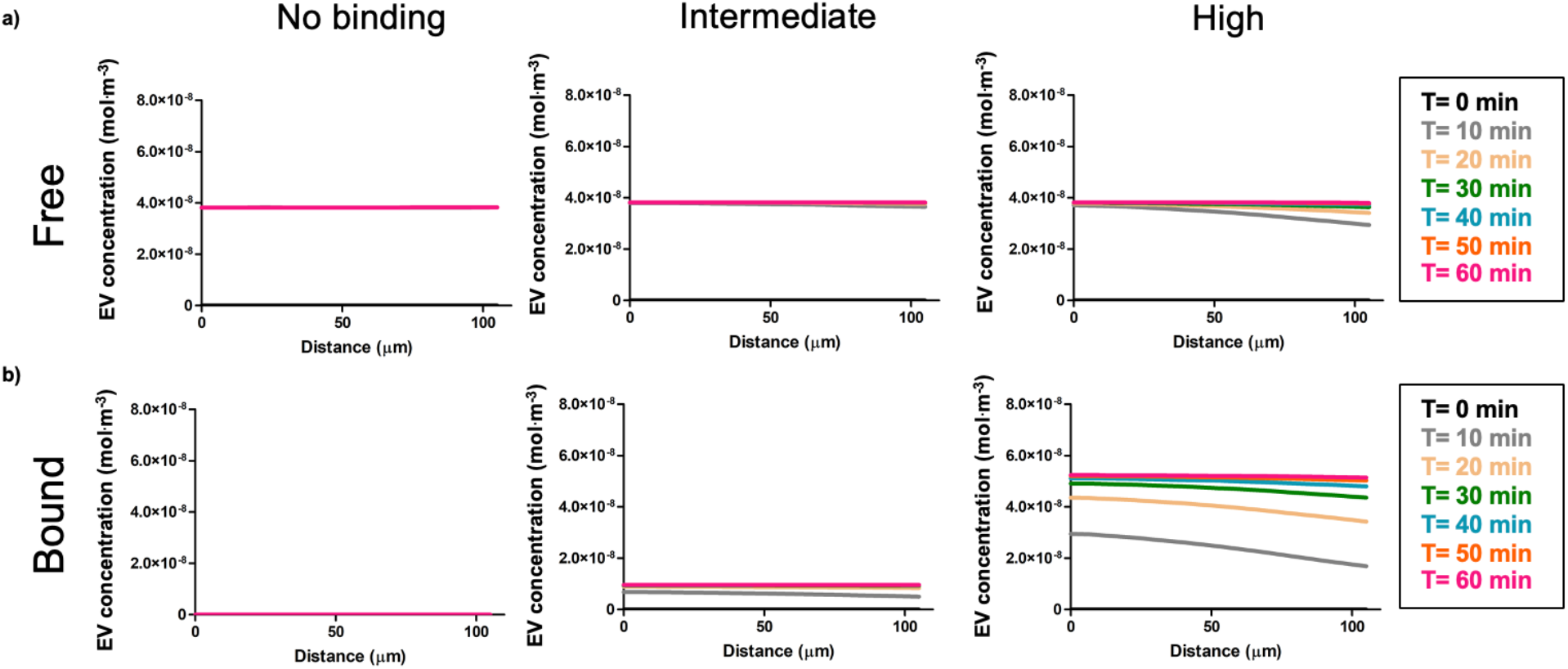
Free and bound EV profiles flatten by T = 60min. a) free and b) bound EV concentration curves flatten over 60 minutes. High affinity EVs reach a higher bound concentration compared to intermediate or no binding EVs.

**Figure S6.**
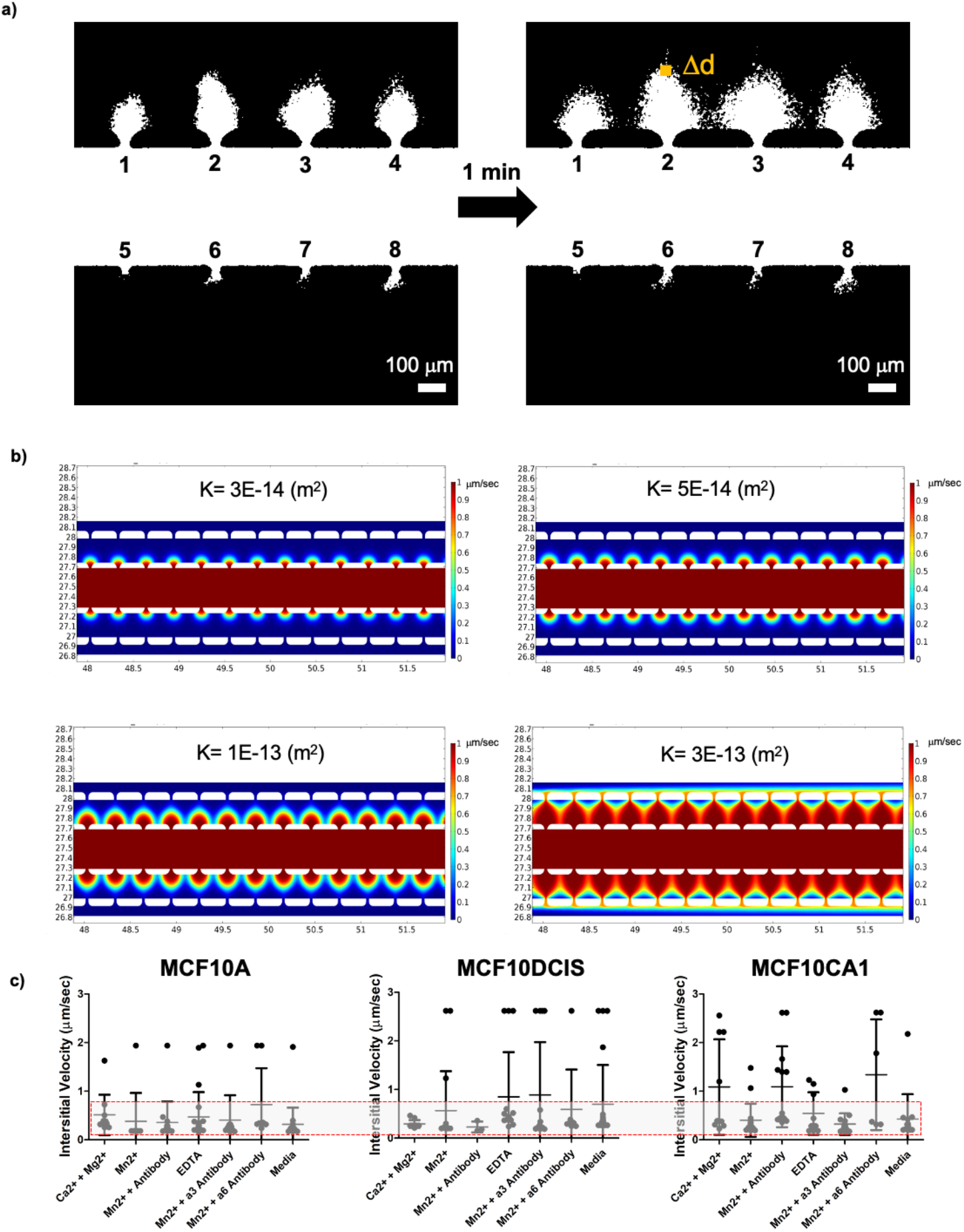
Interstitial fluid velocity was determined via experimental dextran flow and computational modeling. a) Fluorescent dextran profiles were thresholded and resulting representative images demonstrate the distance of dextran transport (Δd) over 1 minute, b) In silico results demonstrate increased dextran transport by increasing the permeability (K) of the ECM. c) Across all convective flow experiments, most interstitial flow velocities were between 0-1 μm/sec. A binned (red lines) interstitial flow velocity range of 0.15-0.75 ym/sec (mean 0.29 μm/sec) was selected for analysis of EV bound concentration.

